# Verification of radical pair mechanism predictions for weak magnetic field effects on superoxide in planarians

**DOI:** 10.1101/2024.11.20.624392

**Authors:** Rishabh, Jana Vučković, Hadi Zadeh-Haghighi, Wendy S. Beane, Christoph Simon

## Abstract

Superoxide concentration and tissue regeneration in planarians exhibit a complex non-monotonic dependence on the strength of an applied weak magnetic field. While this is difficult to understand based on classical physics, a recently proposed quantum model based on a flavin-superoxide radical pair mechanism could replicate the previously observed superoxide concentrations. However, this model also predicts increased superoxide concentrations for both lower and higher fields. This seemed to conflict with earlier experimental observations on blastema sizes, which were correlated with superoxide in the previously observed regime but were known not to follow the predicted trends for lower and higher fields. Motivated by this apparent contradiction, we here directly experimentally tested the predictions of the quantum model for superoxide for lower and higher fields. To our own surprise, our experiments confirmed the predictions of the radical pair model for superoxide, and incorporating interactions with multiple nuclei further improved the model’s agreement with the experimental data. While open questions remain regarding the exact relationship between blastema sizes and superoxide, which is revealed to be more complex than previously observed, and the detailed properties of the underlying radical pair, our results significantly support a quantum biological explanation for the observed magnetic field effects.

## 1 Introduction

Hundreds of studies show that exposure to weak magnetic fields (WMFs), with a magnitude of a few milliTesla (mT) or less, can influence many biological processes, even though the corresponding magnetic energies are much weaker than the thermal energies at room temperature [1]. In particular, researchers have shown in multiple scenarios that the cellular production of reactive oxygen species (ROS) is sensitive to WMFs [2–5]. In many other studies involving WMF effects on higher-level processes, it has been shown that these effects are mediated by modulating ROS concentration [6–9]. ROS are biologically important derivatives of oxygen that are vital for various cellular processes, including signaling [10] and include both free radicals and non-radical species. Superoxide 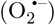 and hydrogen peroxide (H_2_O_2_) are two of the most important members of the ROS family.

Van Huizen et al. conducted a study involving planarian flatworms and found their regeneration to be sensitive to WMFs within the range of 0 − 600 *µ*T [11]. A subsequent study, Kinsey et al., later extended this range to 900 *µT* [12]. Planarians have a large number of somatic stem cells, which account for roughly a quarter of their total cell population [13]. Due to this large adult stem cell population, they have an astonishing capability for regenerating all tissues, including the central nervous system [14]. Van Huizen et al. observed that WMFs altered stem cell proliferation and subsequent differentiation by changing ROS accumulation at the wound site. Although these data established ROS-mediated WMF effects on planarian regeneration, the specific ROS involved remained an open question. In a later study, 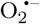, but not H_2_O_2_, was identified as the specific ROS being modulated [12].

Given that, the corresponding energies for WMFs in this range are far smaller than thermal energies at room temperature, no obvious classical explanation is available for these magnetic field effects. However, the radical pair mechanism (RPM) [15, 16] is a potential quantum mechanical explanation for such effects. The RPM involves the simultaneous creation of a pair of radicals, for example, through the transfer of a single electron from one molecule to another. A radical is a molecule that contains at least one unpaired electron. The spins of the two unpaired electrons, one on each constituent molecule of the radical pair (RP), undergo a transient coherent evolution. Depending upon the initial spin configuration of participating molecules, RPs usually start in either singlet or triplet initial states. A system with a total spin equal to 0 (1) has 1 (3) corresponding spin state(s) and is hence termed a singlet (triplet). RPs interact with nearby nuclear spins through hyperfine (HF) interactions and with external magnetic fields via the Zeeman interaction. As neither singlet nor triplet states are stationary states of the spin Hamiltonian, these interactions cause singlet-triplet interconversion. Altering the external magnetic field or substituting an isotope can modify this interconversion. A key feature of the RPM is that the chemical products are spin-selective, with singlet and triplet states leading to different outcomes. As a result, changes in the external magnetic field affect the yields of products formed via the RPM.

In recent years, the RPM has been proposed as an explanation for several WMF effects in biology [1, 17], including several experiments involving WMF effects on ROS production. Usselman et al. proposed a flavin and superoxide-based RPM to explain the effects of oscillating magnetic fields at Zeeman resonance (1.4 MHz) on ROS production in human umbilical vein endothelial cells [18]. A similar mechanism was used to explain the modulation of ROS production in a hypomagnetic environment, which in turn affected neurogenesis in the hippocampal region of mice [19].

In an earlier work, Rishabh et al. studied the possibility of an RPM-based mechanism to explain the effects of WMFs on planarian regeneration [20]. In particular, they investigated the viability of a flavin-superoxide-based radical pair mechanism to explain the observed modulation of 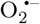 production by WMFs. They found that a triplet-born free radical pair can replicate the previously observed magnetic field dependence for 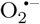. However, some of the predictions of this model seemed to conflict with experimental observations on planarian new tissue growth (blastema size) at hypomagnetic and higher field values (500 − 900 *µ*T). The blastema is a collection of undifferentiated adult stem cell progeny that arises in response to injury and serves as the basis for new tissues during regenerative growth.

Here, we set out to test these predictions of the RPM in planarians for WMF effects on 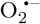 levels. Since data on 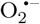 levels were not available for these field strengths, we performed new measurements to test the radical pair model’s predictions for 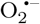 under these conditions. Our experiments confirmed the theoretical predictions for 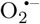 behavior. These results also imply that the interrelationship between 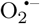 levels and blastema size is nonlinear and more complex than previously thought. Going beyond previous modeling work, we found that incorporating HFIs with multiple nuclei improved the agreement between the theoretical predictions and the experimental observations. Although there remain some open questions regarding an RPMbased modulation of 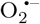 during planarian regeneration, this study significantly supports the possibility of such an underlying mechanism.

## 2 Relevant prior results

### 2.1 Prior Experimental data (Ref. [11, 12])

Van Huizen et al. [11] reported that WMFs alter stem cell proliferation and differentiation, hence regulating blastema formation following amputation in planarians. These effects were dependent on field strength across a wide range, with maximum effects seen at 200 and 500 *µ*T. Significant reductions in blastema size were observed for 200 *µ*T, while a substantial increase was seen at 500 *µ*T.

The observations of Van Huizen et al. [11] also highlighted the importance of ROS, which peak at the wound site 1 hour post injury. They found that pharmacological inhibition with the general ROS inhibitor diphenyleneiodonium resulted in a considerable decrease in blastema size. Moreover, they found that by inhibiting superoxide dismutase (SOD), an enzyme that catalyzes 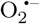 removal, they were able to rescue blastema growth in 200 *µ*T fields. They also found that SOD inhibition significantly increases blastema size in planarians exposed to control geomagnetic conditions. Based on this evidence, they hypothesized that WMF effects were mediated by changing ROS concentrations. To confirm this hypothesis, they measured the ROS levels using a general oxidative stress indicator dye at 1 hour after injury for worms exposed to 200 and 500 *µ*T fields. As expected, measurements at 200 *µ*T revealed a significant decrease in ROS levels, whereas at 500 *µ*T, they saw significantly increased ROS concentrations.

To gain a better understanding of the specific targets of WMFs, Kinsey et al. [12] studied the effects of WMF exposure on 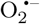 and H_2_O_2_ levels during planarian regeneration. They exposed amputated planarians to 200 and 500 *µ*T and then measured 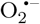 levels using a superoxide specific indicator dye at 1 and 2 hours after injury. 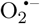 concentrations were found to be sensitive to WMFs in a fashion similar to WMF effects on ROS-mediated stem cell activity. They reported that 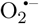 concentration decreased for worms exposed to 200 *µ*T at both 1 and 2 hours post amputation. In contrast, while no significant change was observed for 500 *µ*T fields after 1 hour, a substantial increase was recorded at 2 hours post-amputation. The WMF effects on 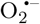 levels for both these field values were significantly greater 2 hours after amputation than 1 hour after. Their 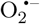 measurements at 2 hours are reproduced in Fig. 2a. It should also be noted that they did not observe any significant changes in H_2_O_2_ concentration as compared to geomagnetic control. Based on these findings, Kinsey et al. concluded that WMF effects on planarian regeneration are mediated at least in part via 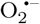.

### 2.2 Theoretical modeling of prior experimental data (Ref. [20])

Can an RPM-based 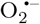 production scheme explain the above observations? As the sign inversion in the product yields at low fields is expected for the RPM, it is not unreasonable to contemplate the existence of such a mechanism [16, 21–23]. To answer the above question, Rishabh et al. [20] compared the predictions of a potential RPM model for O_2_ ^−^ yield with the effects observed by Kinsey et al. for 200 and 500 *µ*T exposures. A detailed summary of the main findings is provided below. Before we proceed, it should be noted that the observational techniques available for superoxide measurement in live planarians provide relative concentrations rather than precise values. Therefore, we will restrict our comparison to the shape of the magnetic field profile rather than the exact values of these effects.

The two primary cellular sources of 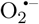 are the mitochondrial electron transport chain and a membrane enzyme family called NADPH oxidase (Nox) [24–27]. In mitochondria, most 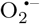 is produced at two sites in complex I and one in complex III. Of the two chemical processes responsible for 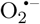 production in mitochondrial complex I, one involves an electron transfer from the reduced form of flavin mononucleotide (FMNH^−^) to molecular oxygen (O_2_) forming 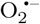 and FMNH [28]. Nox are flavohemoproteins and electron transporters, and Nox1-3 and Nox5 are known to produce 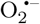. This involves an electron transfer from FADH^−^ to O_2_, occupying a binding site near the heme groups [29]. The electron transfers from fully reduced flavin to O_2_ during the production of 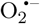 in both mitochondria and Nox, as well as the magnetic field dependence of 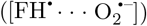 production, suggest the involvement of a flavin-superoxide RP 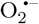. This FH and 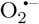 based RPM can serve as the basis for explaining various WMF effects observed in the context of 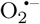 production [6, 12].

Following Usselman et. al.[18], a triplet-born RP system of FH^·^ and 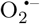 (See Fig. 1a) was proposed with triplet and singlet products being 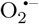 and H_2_O_2_, respectively (See Fig. 1b). The singlet and triplet reaction rates are denoted by *k*_*S*_ and *k*_*T*_, respectively. The spin relaxation rates of radicals A and B are denoted by *r*_*A*_ and *r*_*B*_, respectively.

**Figure 1:**
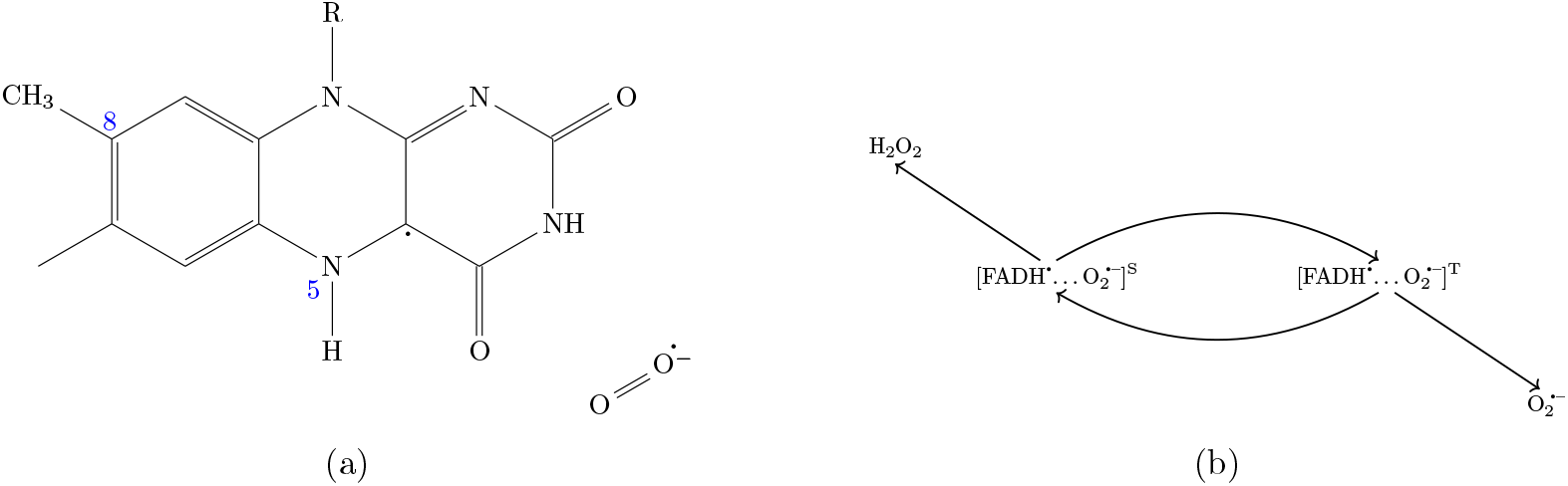
Radical pair model: (a) Flavin-superoxide radical pair. (b) Radical pair reaction scheme. Triplet and singlet products are 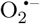 and H_2_O_2_, respectively. In this study we consider the following five isotropic HF couplings for FH : H5 (−802.9 *µ*T), N5 (431.3 *µ*T), three H8 (255.4 *µ*T) [30].

To study the RP dynamics, a simplified spin Hamiltonian including only the Zeeman and largest isotropic HF coupling for FH^·^ was considered:

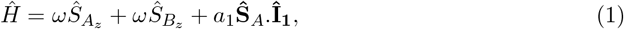

where 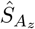 and 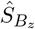 are the spin-z operators of radical electron *A* (FH^·^) and 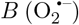, respectively, *ω* is the Larmor precession frequency of the electrons due to the Zeeman effect, 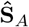 is the spin vector operator of radical electron *A*, **Î**_**1**_ is the nuclear spin vector operator of the H5 of FH^·^, and *a*_1_ is the isotropic HF coupling constant (HFCC) between the H5 of FH^·^ and the radical electron A (*a*_1_ = −802.9 *µ*T) [30]. H5 has by far the largest isotropic HFCC among all the nuclei in FH [30].

The fractional triplet 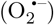 yield generated by the RPM can be determined by monitoring the dynamics of RP spin states. For details of the calculations, see the Methods section. The ultimate fractional triplet yield 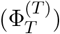 for an RP that originates in a triplet state, when considering time intervals significantly longer than the RP’s lifetime, is as follows:

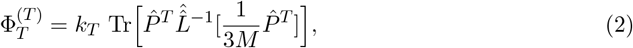

where 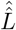 is the Liouvillian superoperator, 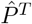 is the triplet projection operator, *M* is the total number of nuclear spin configurations, and *k*_*T*_ is the triplet reaction rate.

There are four free parameters in this model, namely, *k*_*S*_, *k*_*T*_, *r*_*A*_, and *r*_*B*_. The question was whether there are regions in parameter space where the simulated behavior corresponds to the experimental observation (i.e., a positive change in 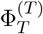 at 500 *µ*T and a negative change at 200 *µ*T with respect to geomagnetic control). For this purpose, Rishabh et al. investigated the signs of triplet yield changes with respect to control at 200 *µ*T and 500 *µ*T over a wide range of chemical reaction rates (*k*_*S*_ ∈ {10^4^ *s*^−1^, 10^7^ *s*^−1^} and *k*_*T*_ ∈ {10^4^ *s*^−1^, 10^7^ *s*^−1^}) for various pairs of spin relaxation rates *r*_*A*_ and *r*_*B*_. They observed that such a region in *k*_*S*_-*k*_*T*_ plane can be found provided *r*_*A*_ ≤ 10^5^ *s*^−1^ and *r*_*B*_ ≤ 10^6^ *s*^−1^.

Fig. 2b shows the change in 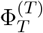 with respect to the geomagnetic control as a function of the magnetic field for various values of *k*_*S*_ and *k*_*T*_ when *r*_*A*_ is fixed at 1 × 10^5^ *s*^−1^ and *r*_*B*_ is fixed to 1 × 10^6^ *s*^−1^. It is clear that for appropriate rate values, Rishabh et al.’s RP model can replicate the previously observed magnetic field dependence, including the sign change. However, it should be noted that the magnitude of the observed effects is much larger than what could be achieved by any RP model. This highlights the necessity of an amplification mechanism, as discussed by Rishabh et al. We will revisit this issue in the discussion section.

**Figure 2:**
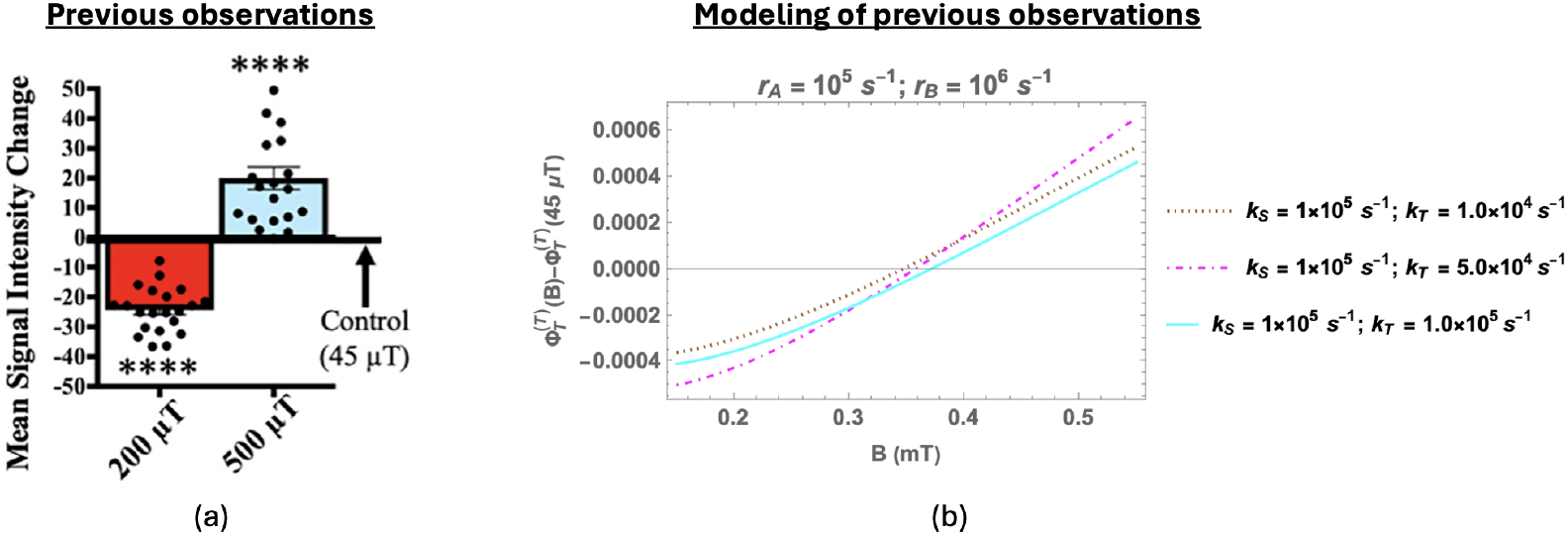
Summary of relevant prior results: (a) Superoxide 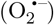 measurements at the wound site 2 hours after amputation (reproduced from [12]). 200 (red) and 500 (blue) *µ*T exposures each relative to 45 *µ*T geomagnetic controls. Significance: Student’s t-test. **** p*<*0.0001. (b) Change in the fractional triplet yield for triplet-born flavin-superoxide RP with respect to the geomagnetic control (45 *µ*T) as a function of the magnetic field (150 − 550 *µ*T) [20]. Here, only the largest isotropic HF coupling (H5) is taken into account (HFCC value is −802.9 *µ*T). *r*_*A*_ = 10^5^ *s*^−1^ and *r*_*B*_ = 10^6^ *s*^−1^. *k*_*S*_ and *k*_*T*_ are singlet and triplet reaction rates, respectively. *r*_*A*_ and *r*_*B*_ are the spin relaxation rates of radicals A and B, respectively.

### 2.3 Predictions of the radical pair model

#### Beyond 500 *µ*T

As shown in Fig. 3 (yellow shaded region), the RPM predicts that as we increase the magnetic field strength beyond 500 *µ*T, we should observe a corresponding rise in 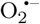 levels. Note that, the exact amount of this rise will depend on the specific parameters of the model.

**Figure 3:**
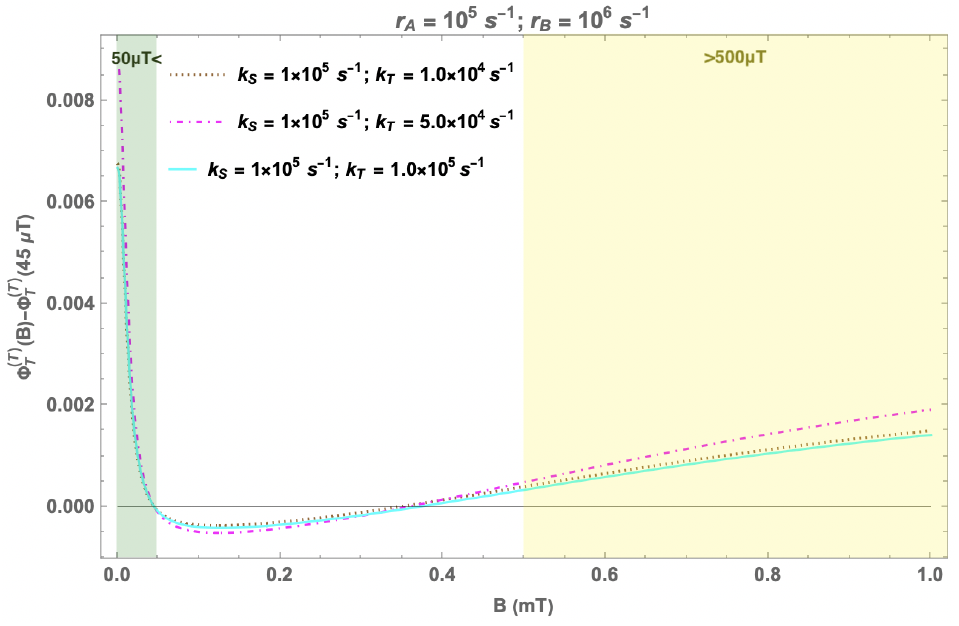
Predictions of the RPM model: Change in the fractional triplet yield for triplet-born flavin-superoxide RP with respect to the geomagnetic control (45 *µ*T) as a function of the magnetic field (0 − 1 mT). *r*_*A*_ = 10^5^ *s*^−1^ and *r*_*B*_ = 10^6^ *s*^−1^. *k*_*S*_ and *k*_*T*_ are singlet and triplet reaction rates, respectively. *r*_*A*_ and *r*_*B*_ are the spin relaxation rates of radicals A and B, respectively. Again, only the largest isotropic HF coupling (H5) is taken into account (HFCC value is −802.9 *µ*T). The region with sub-geomagnetic fields is shaded in green, and the region with fields greater than 500 *µ*T is shaded in yellow.

#### Hypomagnetic effects

In an RPM, the fractional triplet yield can also be altered by shielding the geomagnetic field [31]. The effect on the fractional triplet yield of the RPM of shielding geomagnetic field for a tripletborn RP is shown in Fig. 3 (green shaded region). These simulations suggest a positive change in a hypomagnetic environment. The exact size of the effect will again depend on the specific parameters of the model [19, 31].

## 3 Results

### 3.1 Measurement of magnetic field effects on superoxide

Despite the fact that the predictions of the flavin-superoxide RP model can align with the observed behavior of 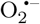 at 200 and 500 *µ*T, its predictions at the hypomagnetic and higher fields might raise serious doubts about the viability of the model. Kinsey et al. measured the effects of hypomagnetic and higher fields (600-900 *µ*T) on blastema size and general ROS, but not specifically on 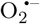. Their observations showed no significant effects (except at 900 *µ*T) for these fields. If similar patterns are reflected in 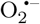 levels, it would challenge the current model unless some deamplification mechanism is activated (or the amplification mechanism is deactivated) when 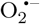 levels become too high. According to the existing hypothesis regarding 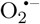 mediation of WMF effects on planarian regeneration, it is expected that 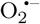 levels should follow a behavior similar to that of blastema size. Therefore, to settle the question of the involvement of the RPM, we conducted measurements of 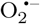 levels at 700 and 900 *µ*T, as well as at hypomagnetic field values. To our surprise, the experiments confirmed the theoretical predictions of the RP model concerning the behavior of superoxide at hypomagnetic and larger fields, contrary to expectations based on earlier experimental observations on blastema sizes.

Fig. 4 shows the results of these experiments. Adult *Schmidtea mediterranea* planarians were amputated above the pharynx (feeding tube) to produce fragments that undergo head regeneration as illustrated in Fig. 4a. Regenerates were exposed to static WMFs at 0, 200, 500, 700 and 900 *µ*T for the first 2 hours post amputation. The superoxide-specific live reporter dye, orange 1, was used to visualize 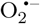 concentrations at the wound site at 2 hours post injury (the peak of superoxide accumulation) as in Fig. 4b. Quantification of these data is shown in Fig. 4c, demonstrating that similar to previous findings [12], 200 *µ*T significantly inhibited while 500 *µ*T significantly increased superoxide levels as compared to geomagnetic controls. However, in contrast to those investigations of general ROS and blastema size, examination revealed that 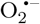 concentrations are also significantly increased at 0, 700, and 900 *µ*T. Furthermore, the peak average increase in 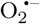 occurred with 0 *µ*T exposure, as predicted by the RP model. Together, these experimental results demonstrate that WMFs alter superoxide concentration at the wound site in a field strength dependent manner consistent with the RP model.

**Figure 4:**
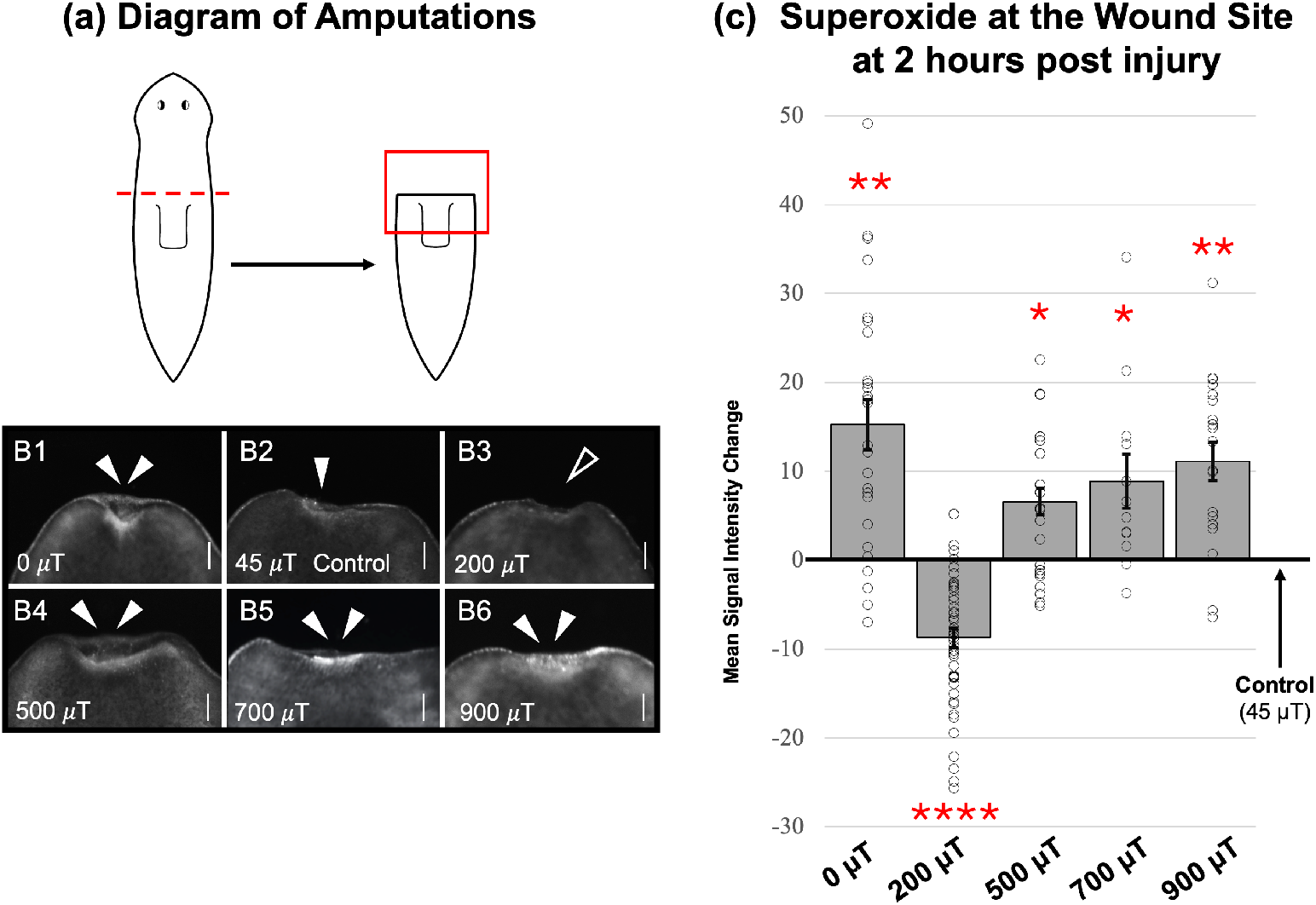
Validation of RPM predictions: Regenerating *S. mediterranea* planarians 2 hours after injury, exposed to a range of static WMFs. (a) Diagram of amputation (dotted red line). Red box represents region as shown in b which corresponds to the anterior wound site. (b) 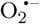 accumulation visualized by orange 1 live reporter dye. Control = 45 *µ*T (B2, geomagnetic average). Solid arrow = normal 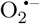 levels. Double arrows = increased and open arrow = inhibited O_2_ ^−^ levels (as compared to control). Scale bars = 100 *µ*m. Anterior is up. (c) Quantification of b, with changes in signal intensity relative to 45*µ*T controls. n*>*12 for all. Significance: Student’s t-test. *p*<*0.02, **p*<*0.005, ****p*<*0.0001.

### 3.2 Multiple hyperfine interactions

Although Rishabh et al.’s model correctly predicted the sign of WMF effects, it significantly overestimates the impact of hypomagnetic fields compared to higher field values. We found that this is, in part, an artifact of the simplifying assumption of including only one HFI. Bringing the RP model closer to reality by taking into account isotropic HFIs with multiple nuclei (not just the largest one as in the previous work by Rishabh et al.) leads to a much-improved correspondence between the predictions of the theoretical model and the observations from the experiments. Fig. 5 shows the theoretical predictions of our model with five HFIs. We have taken into account the five nuclei with the largest isotropic HFCCs, namely: H5 (−802.9 *µ*T), N5 (431.3 *µ*T), three H8 (255.4 *µ*T) [30]. Note that introducing a second HFI had significant effects, but adding additional HFIs beyond that had little impact. This is shown in Fig. 6 in the supporting information. At this point, let us also note that, despite the introduction of multiple HFIs, the agreement between theory and experiment—though significantly improved—is still not perfect. While it is difficult to pinpoint the exact cause of this mismatch, it may stem from the amplification chemistry or the observational techniques used for measuring superoxide.

**Figure 5:**
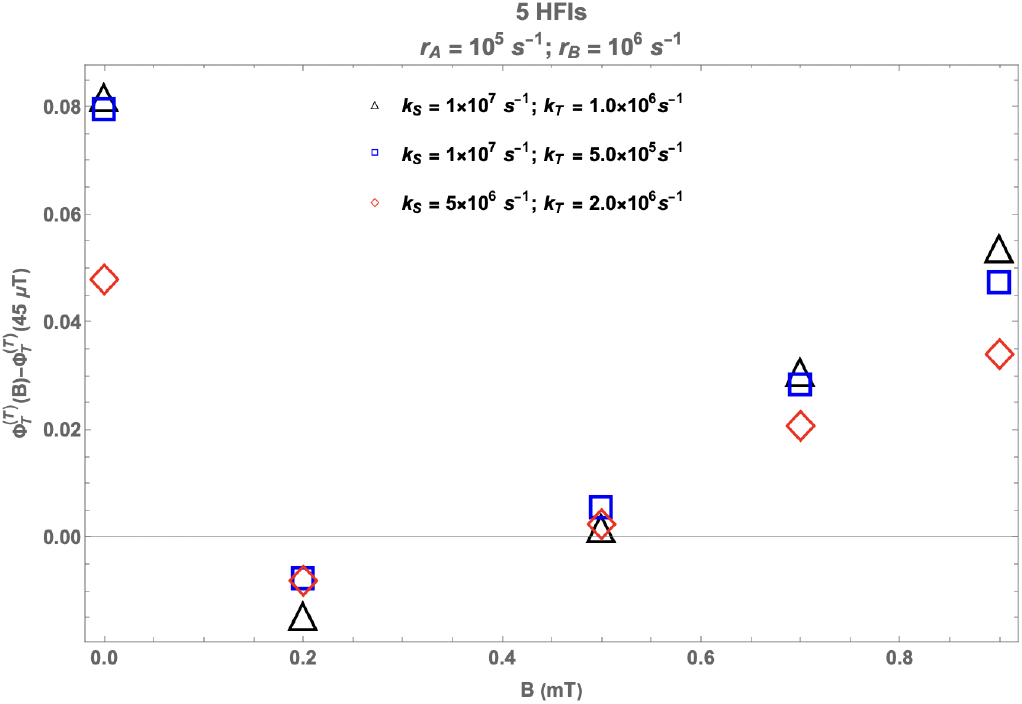
Multiple HFIs: Change in the fractional triplet yield for triplet-born RP with respect to the geomagnetic control (45 *µ*T) as a function of the magnetic field. *r*_*A*_ = 10^5^ *s*^−1^ and *r*_*B*_ = 10^6^ *s*^−1^. *k*_*S*_ and *k*_*T*_ are singlet and triplet reaction rates, respectively. *r*_*A*_ and *r*_*B*_ are the spin relaxation rates of radicals A and B, respectively. 5 HFIs: H5 (−802.9 *µ*T), N5 (431.3 *µ*T), three H8 (255.4 *µ*T) [30].

**Figure 6:**
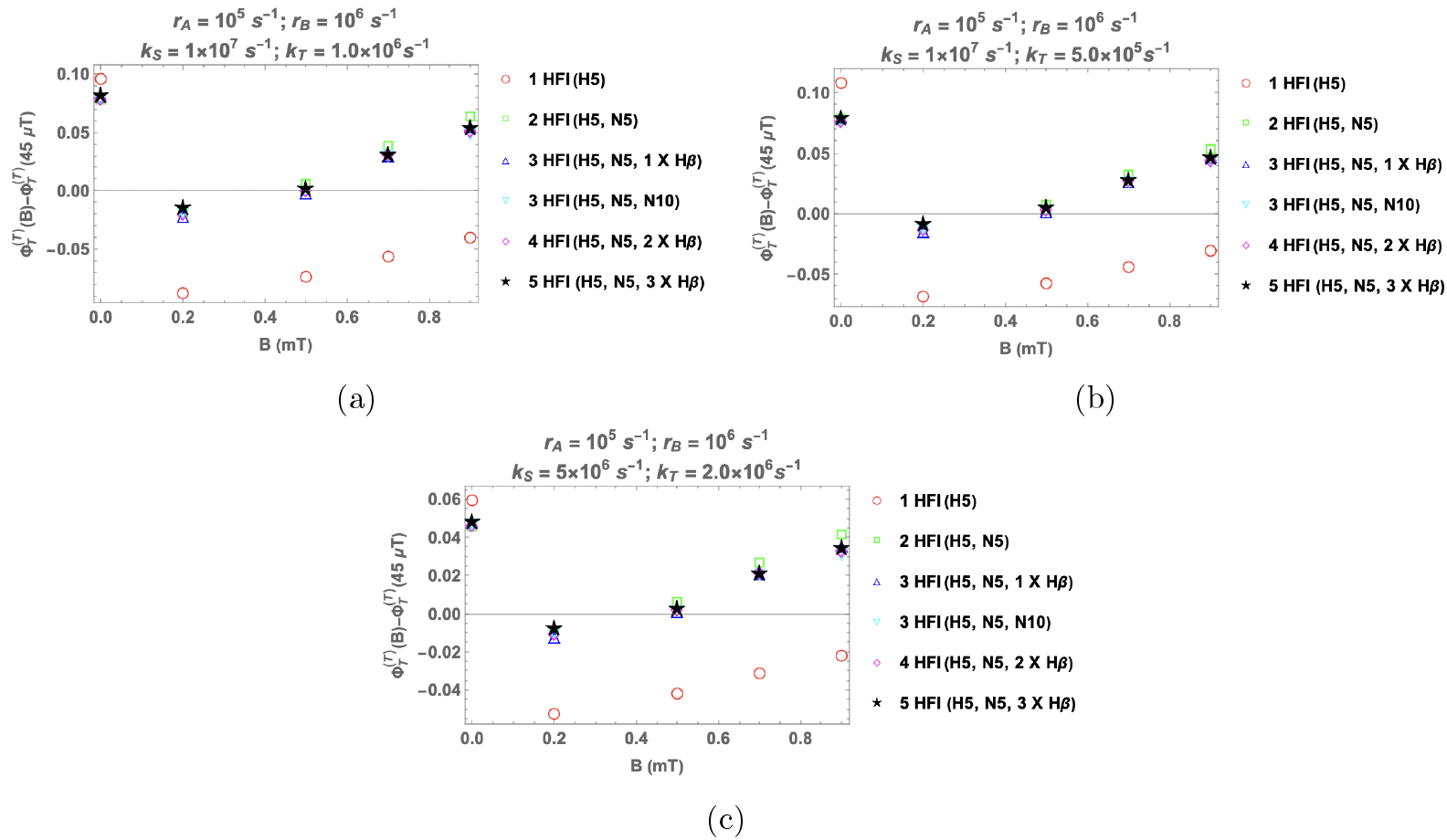
Effects of having more than one HFIs: Change in the fractional triplet yield for triplet-born RP with respect to the geomagnetic control (45 *µ*T) as a function of the magnetic field. *r*_*A*_ = 10^5^ *s*^−1^, *r*_*B*_ = 10^6^ *s*^−1^. *r*_*A*_ and *r*_*B*_ are the spin relaxation rates of radicals A and B, respectively. *k*_*S*_ and *k*_*T*_ are singlet and triplet reaction rates, respectively.(a) *k*_*S*_ = 10^7^ *s*^−1^ and *k*_*T*_ = 10^6^ *s*^−1^, (b) *k*_*S*_ = 10^7^ *s*^−1^ and *k*_*T*_ = 5 × 10^5^ *s*^−1^, (c) *k*_*S*_ = 5 × 10^6^ *s*^−1^ and *k*_*T*_ = 2 × 10^6^ *s*^−1^

## 4 Discussion

In this work, we set out to test the predictions of a 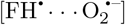 RP-based model for WMF effects on 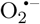 levels during planarian regeneration. It was known that a triplet-born free radical pair can replicate the previously observed magnetic field dependence, including the sign change [20]. However, the model’s predictions at hypomagnetic and higher fields did not align with the expected behavior of 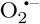 based on prior observations of blastema size. Surprisingly, our experiments confirmed the predictions of the radical pair model concerning the behavior of superoxide at hypomagnetic and larger fields. Moreover, extending previous models, we found that taking into account isotropic HFIs with multiple nuclei leads to a much-improved correspondence between the RP model’s predictions and the experimental data. These results strongly suggest the possibility of an underlying RPM.

These results also highlight the complex interrelation between 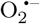 and tissue regeneration in planarians. As mentioned above, the blastema size, measured by Kinsey et al. [12], does not emulate the behavior of superoxide concentration at the wound site, in particular for 0 and 700 *µ*T. This non-linear relationship between new tissue growth and superoxide levels after injury may be related to the fact that superoxide accumulation occurs in the first hours after injury, while blastema growth occurs between 24-72 hours [32]. Teasing apart the exact relationship between early ROS and tissue regrowth should be a focus of studies going forward.

It should be noted that despite the successful predictions of this RP model regarding the superoxide levels, some open questions remain. We highlight some of the main issues in this and the following paragraphs. The usual singlet product of the 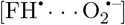 RP is H_2_O_2_ [18, 19]. However, Kinsey et al.[12] did not observe any significant effect of WMF on H_2_O_2_ concentration. This suggests that either H_2_O_2_ is not the main singlet product in the present case, or more probably, it indicates the absence of an amplification process for H_2_O_2_.

It has been suggested in the past that due to fast molecular rotation, free 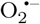 should have a spin relaxation lifetime on the orders of 1 ns and hence a fast spin relaxation rate *r*_*B*_ [33, 34]. The relaxation rate requirement calculated by our model for *r*_*B*_ is significantly lower than this expected value. However, this fast spin relaxation of free superoxide can be lowered if the molecular symmetry is reduced and the angular momentum is quenched by the biological environment [33, 34]. Although, it should be noted that for this to happen 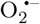 must be tightly bound [34]. Such a possibility may arise in the case of the Nox enzyme because of the presence of O_2_ binding pockets near the heme proteins. However, it should be noted that tightly bound flavin molecules, which require consideration of anisotropic rather than isotropic hyperfine coupling, cannot explain experimental observations [20]. This strongly suggests that the 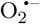 involved is not produced via Nox, or the flavin bound to Nox is unexpectedly still relatively free to rotate. It has also been indicated that O_2_ would need to bind in the mitochondrial electron transfer flavoprotein for superoxide production [35]. Direct evidence of such inhibition of spin relaxation (for example, an electron paramagnetic resonance spectrum of 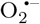) has yet to be found.

Despite predicting the correct behavior of magnetic field effects, the RPM model alone can not predict the right size of these effects and does not account for the temporal aspect of Kinsey et al.’s [12] observation. This illustrates the need for an amplification process for 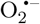 [20]. The existence of 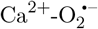 self-amplifying loop [36] and JNK-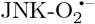 amplification pathway [37], and the fact that such a pathway is activated precisely during regeneration [38] adds to the plausibility of such an amplification process.

It should also be pointed out that we have ignored inter-radical interactions in our modeling. The effects of including exchange interaction have been studied in Ref. [20] and do not change our main conclusions.

In this study, we have only considered triplet-born free RPs. However, other related possibilities, such as F-pairs and radical triads, cannot be ruled out [20]. Moreover, the possibility that these WMF effects may be due to some other RP, such as flavin-tryptophan, can not be completely excluded. The production of 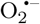, in that case, might happen downstream of the RP spin dynamics [39]. However, it should be noted that there is no strong biological reason to believe the involvement of such RPs in 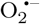 production during planarian regeneration. For example, it remains unclear whether CRY, a natural host of flavin-tryptophan RP, plays any role in planarians. It is also possible that mechanisms other than the RPM could also explain the WMF effects on planarians.

In summary, although further investigation is needed to conclusively prove the involvement of a radical pair in planarian regeneration or to determine the exact nature of such a pair, the experimental verification of RPM’s predictions regarding superoxide levels in this study provides significant support to the possibility of such an underlying quantum mechanism.

## 5 Methods

### 5.1 Radical pair mechanism calculations

The state of the RP is described using the spin density operator. The coherent spin dynamics, chemical reactivity, and spin relaxation all together determine the time evolution of the spin density matrix of the RP system.

Since the ground state of the oxygen molecule is a triplet, we will consider the initial state of the RP to be a triplet:

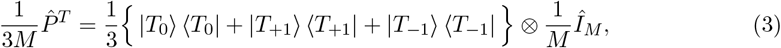

where 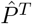 is the triplet projection operator, *M* is the total number of nuclear spin configurations, |*T*_0_⟩ and |*T*_*±*1_⟩ represent the triplet states of two electrons in RP with the spin magnetic quantum number (*m*_*S*_) equal to 0 and ±1 respectively. *Î*_*M*_ represents the completely mixed initial state of the nuclei.

The time dependence of the spin density operator is obtained using the Liouville Master Equation [16, 40]:

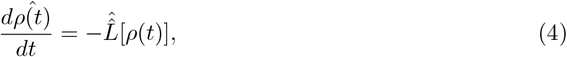

where Liouvillian superoperator 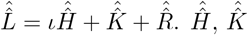, and 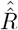 are Hamiltonian superoperator, chemical reaction superoperator, and spin relaxation superoperator, respectively.

The most general spin Hamiltonian for RP will include Zeeman 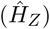 and HF 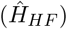 interactions as well as the inter-radical interactions (*Ĥ* _*IR*_), which incorporate exchange and dipolar terms.

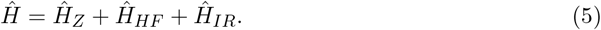

Due to the potential random orientation of the molecules in question, we only take into account the isotropic Fermi contact contributions in HF interactions. In this study we consider the following five isotropic HF couplings for FH ^·^:

**Table 1:** Hyperfine interactions taken into account for FH^·^ [30].

| Nuclei | HFCC (μT) |
| --- | --- |
| H5 | -802.9 |
| N5 | 431.3 |
| H8 (X3) | 255.4 |

The unpaired electron on 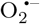 (containing two ^16^O nuclei) has no HF interactions. It should be noted that the fact that 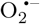 has no HFI helps in improving the magnetic sensitivity of RPs [41–44]. Furthermore, for simplicity, we do not consider any inter-radical interactions in our model. The form of the simplified spin Hamiltonian is given in Eq. 1

For spin-selective chemical reactions (reaction scheme of Fig. 1b), we use the Haberkorn superoperator [40], which is given by the following equation:

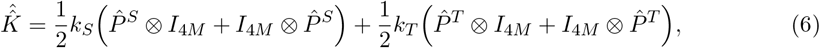

where symbols have above stated meanings. Spin relaxation is modeled via random time-dependent local fields [45, 46], and the corresponding superoperator reads as follows:

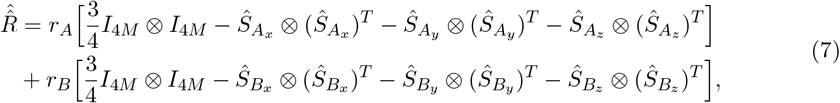

where the symbols have above stated meanings. The ultimate fractional 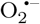 yield for tripletborn 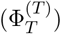 for time periods much greater than the RP lifetime is given by:

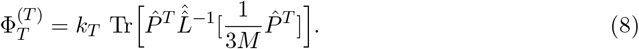

The computational calculations and plotting were performed on Mathematica [47].

### 5.2 Animal care and amputations

An asexual clonal line of *Schmidtea mediterranea* (CIW4) was maintained in the dark at 18 °C. Planarians were kept in Ultrapure Type 1 water with Instant Ocean salts at 0.5 g/L (worm water). Animals were fed every third week with liver paste processed from a whole calf liver (antibiotic and hormone free) obtained from C. Roy & Sons Processing (Yale, MI). Liver paste was frozen and thawed only once. Worms 5-6 mm in length were used (which had been starved at least 2 weeks before use). Amputations were done with a scalpel over a custom made cooling peltier plate under a dissecting microscope. Fragments were produced via transverse amputation just anterior to the pharynx, with a single cut made at 90 degrees to the sagittal plane.

### 5.3 Magnetic field exposure

Experimentally-controlled static WMF exposures were generated with a custom-built MagShield box (a pair of triaxial Helmholtz coils inside a partitioned mu-metal enclosure that blocks external magnetic fields), as previously described [48]. Direct electric current to Helmholtz coils was supplied by DC power sources (Mastech HY3005D-2-R) and was fed through both x and y axis coils to produce a uniform magnetic field. The MagShield box was kept in a temperature-controlled room (20 °C) and experiments were performed in the dark. Animals were placed in 60 mm Petri dishes in worm water (or specific media as described), with a max of n = 10 per replicate. For each replicate, 45 *µ*T (Earth normal) controls were run in one partition concurrently with experimental field strengths (as indicated) in the other partition. Field strengths were confirmed using a milli/Gauss meter (AlphaLab models GM1-HS or MGM) at the start and completion of each run. All planarians were exposed to WMFs within 5 min of amputation and then continuously (except when dye solution was added) until they were removed for imaging. Replicates (N) and total samples (n) per condition: 45 *µ*T N = 15, n = 97; 0 *µ*T N = 3, n = 26; 200 *µ*T N = 6, n = 48; 500 *µ*T N = 3, n = 29; 700 *µ*T N = 2, n = 12; 900 *µ*T N = 2, n = 20. Note: 0 *µ*T = +/-2 *µ*T (tolerance of milligauss meter).

### 5.4 Detection of superoxide and statistical analyses

Superoxide levels were detected using a cell-permeant live fluorescent reporter dye as previously described [12]. 2 *µ*M orange 1 dye (Enzo Life Sciences ENZ-51012) in worm water used used, made from 5 mM dimethylformamide stock; excitation, 550 nm; emission, 620 nm. Fragments were exposed to WMFs from 5 min to 1 h post amputation. At 1 h, fragments were quickly moved to new 35 mm Petri dishes in orange 1 solution and returned to the MagSheild box for an additional 1 h of WMF exposure. Thus fragments were exposed to WMFs for 2 h total, including 1 h of dye loading, at which time regenerates were rinsed 3X in ice cold worm water in the dark to preserve fluorescence and imaged. A Zeiss V20 fluorescence stereomicrope with an AxioCam MRm camera and ZEN (lite) software was used for image collection. Live images were taken while fragments were extended to prevent signal intensity skewing due to scrunching. Animals were imaged in 35 mm FluoroDishes (WPI FD35-100) with 25 mm round no. 1.5 coverslips (WPI 503508). All samples were imaged at the same magnification and exposure levels to prevent confounding variables during comparisons (i.e., acquisition conditions were kept constant across an experiment between control/treated). Photoshop (Adobe) was used to orient and scale images. No data was added or subtracted. Original images available by request. For quantification: the magnetic lasso tool in Photoshop was used to measure gray mean values (signal intensity) of fluorescent dye at the anterior wound. To account for any variation in dye loading, signal intensity was calculated as the difference between signal at the anterior wound site versus signal from the middle of the regenerate (the pharyngeal region): blastema – pharyngeal region. Significance: two-tailed Student’s t-test with unequal variance (Microsoft Excel) as compared to Earth normal controls.

## Data availability

The original contributions presented in the study are included in the article/Supplementary Material, further inquiries can be directed to the corresponding author.

## Code availability

The Mathematica notebooks used to generate theoretical plots are available from the corresponding author upon request.

## Acknowledgement

The authors thank Dr. Luke Kinsey and Prof. Dennis Salahub for helpful discussions. This work involved the use of Advanced Research Computing (ARC) cluster at the University of Calgary.

This work was supported by the Natural Sciences and Engineering Research Council through its Discovery Grant and CREATE programs as well as the Alliance Quantum Consortia Grant ‘Quantum Enhanced Sensing and Imaging’, and by the National Research Council of Canada through its Quantum Sensing Challenge Program. This work was further supported by National Science Foundation grant NSF-2105474 and National Institutes of health grant NIH-1R15GM150073-01 to W.S.B. Funding was also provided to J.V. by the Fulbright Foreign Student Program, which is sponsored by the U.S. Department of State.

## Author contributions

CS and WSB conceived the project; R performed the theoretical modelling and calculations with help from HZH and CS; JV performed planarian experiments; WSB and JV analyzed the planarian data; R and JV wrote the original draft with feedback from HZH, WSB and CS; R, JV, HZH, WSB and CS reviewed and edited the final version.

## Supporting Information

### Multiple hyperfine interaction

We found that introducing a second HFI had significant effects, but adding additional HFIs beyond that had little impact.

**Table 2:** Hyperfine interactions for FH ^·^ [30].

| Nuclei | HFCC ($\mu\text{T}$) |
| --- | --- |
| H5 | -802.9 |
| N5 | 431.3 |
| H8 (X3) | 255.4 |
| N10 | 250.6 |
| Hβ | 190.8 |

## References

1. Zadeh-Haghighi, H. & Simon, C. Magnetic field effects in biology from the perspective of the radical pair mechanism. Journal of the Royal Society Interface 19, 20220325 (2022).

2. Wang, H. & Zhang, X. Magnetic fields and reactive oxygen species. International Journal of Molecular Sciences 18, 2175 (2017).

3. Calabrò, E. et al. Effects of low intensity static magnetic field on FTIR spectra and ROS production in SH-SY5Y neuronal-like cells. Bioelectromagnetics 34, 618–629 (2013).

4. Martino, C. F. & Castello, P. R. Modulation of hydrogen peroxide production in cellular systems by low level magnetic fields. PLoS One 6, e22753 (2011).

5. Poniedzialek, B., Rzymski, P., Karczewski, J., Jaroszyk, F. & Wiktorowicz, K. Reactive oxygen species (ROS) production in human peripheral blood neutrophils exposed in vitro to static magnetic field. Electromagnetic Biology and Medicine 32, 560–568 (2013).

6. Zhang, B. et al. Long-term exposure to a hypomagnetic field attenuates adult hippocampal neurogenesis and cognition. Nature communications 12, 1174 (2021).

7. Bekhite, M. M. et al. Static electromagnetic fields induce vasculogenesis and chondro-osteogenesis of mouse embryonic stem cells by reactive oxygen species-mediated up-regulation of vascular endothelial growth factor. Stem Cells and Development 19, 731–743 (2010).

8. De Nicola, M. et al. Magnetic fields protect from apoptosis via redox alteration. Annals of the New York Academy of Sciences 1090, 59–68 (2006).

9. Hajipour Verdom, B., Abdolmaleki, P. & Behmanesh, M. The static magnetic field remotely boosts the efficiency of doxorubicin through modulating ROS behaviors. Scientific reports 8, 990 (2018).

10. Sies, H. & Jones, D. P. Reactive oxygen species (ROS) as pleiotropic physiological signalling agents. Nature reviews Molecular cell biology 21, 363–383 (2020).

11. Van Huizen, A. V. et al. Weak magnetic fields alter stem cell–mediated growth. Science advances 5, eaau7201 (2019).

12. Kinsey, L. J., Van Huizen, A. V. & Beane, W. S. Weak magnetic fields modulate superoxide to control planarian regeneration. Frontiers in Physics 10, 1356 (2023).

13. Baguñà, J. The planarian neoblast: the rambling history of its origin and some current black boxes. International Journal of Developmental Biology 56, 19–37 (2012).

14. Cebrià, F. Regenerating the central nervous system: how easy for planarians! Development genes and evolution 217, 733–748 (2007).

15. Closs, G. L. Mechanism explaining nuclear spin polarizations in radical combination reactions. Journal of the American Chemical Society 91, 4552–4554 (1969).

16. Steiner, U. E. & Ulrich, T. Magnetic field effects in chemical kinetics and related phenomena. Chemical Reviews 89, 51–147 (1989).

17. Hore, P. J. & Mouritsen, H. The radical-pair mechanism of magnetoreception. Annual review of biophysics 45, 299–344 (2016).

18. Usselman, R. J. et al. The quantum biology of reactive oxygen species partitioning impacts cellular bioenergetics. Scientific reports 6, 38543 (2016).

19. Rishabh, R., Zadeh-Haghighi, H., Salahub, D. & Simon, C. Radical pairs may explain reactive oxygen species-mediated effects of hypomagnetic field on neurogenesis. PLOS Computational Biology 18, e1010198 (2022).

20. Rishabh Zadeh-Haghighi, H. & Simon, C. Radical pairs and superoxide amplification can explain magnetic field effects on planarian regeneration. arXiv preprint arXiv:2312.06597 (2023).

21. Timmel, C. R., Till, U., Brocklehurst, B., Mclauchlan, K. A. & Hore, P. J. Effects of weak magnetic fields on free radical recombination reactions. Molecular Physics 95, 71–89 (1998).

22. Brocklehurst, B. Magnetic fields and radical reactions: recent developments and their role in nature. Chemical Society Reviews 31, 301–311 (2002).

23. Lewis, A. M. et al. On the low magnetic field effect in radical pair reactions. The Journal of Chemical Physics 149 (2018).

24. Bedard, K. & Krause, K.-H. The NOX family of ROS-generating NADPH oxidases: physiology and pathophysiology. Physiological reviews 87, 245–313 (2007).

25. Terzi, A. & Suter, D. M. The role of NADPH oxidases in neuronal development. Free Radical Biology and Medicine 154, 33–47 (2020).

26. Wallace, D. C., Fan, W. & Procaccio, V. Mitochondrial energetics and therapeutics. Annual Review of Pathology: Mechanisms of Disease 5, 297–348 (2010).

27. Zhao, R.-Z., Jiang, S., Zhang, L. & Yu, Z.-B. Mitochondrial electron transport chain, ROS generation and uncoupling. International journal of molecular medicine 44, 3–15 (2019).

28. Markevich, N. I. & Hoek, J. B. Computational modeling analysis of mitochondrial superoxide production under varying substrate conditions and upon inhibition of different segments of the electron transport chain. Biochimica et Biophysica Acta (BBA)-Bioenergetics 1847, 656–679 (2015).

29. Wu, X. et al. Mechanistic insights on heme-to-heme transmembrane electron transfer within NADPH oxydases from atomistic simulations. Frontiers in Chemistry 9, 650651 (2021).

30. Lee, A. A. et al. Alternative radical pairs for cryptochrome-based magnetoreception. Journal of The Royal Society Interface 11, 20131063 (2014).

31. Zadeh-Haghighi, H., Rishabh, R. & Simon, C. Hypomagnetic field effects as a potential avenue for testing the radical pair mechanism in biology. Frontiers in Physics 11, 1026460 (2023).

32. Birkholz, T. R., Van Huizen, A. V. & Beane, W. S. Staying in shape: Planarians as a model for understanding regenerative morphology. Seminars in Cell Developmental Biology 87. Planarian regeneration, 105–115. issn: 1084-9521. https://www.sciencedirect.com/science/article/pii/S1084952117302057 (2019).

33. Hogben, H. J., Efimova, O., Wagner-Rundell, N., Timmel, C. R. & Hore, P. Possible involvement of superoxide and dioxygen with cryptochrome in avian magnetoreception: origin of Zeeman resonances observed by in vivo EPR spectroscopy. Chemical Physics Letters 480, 118–122 (2009).

34. Player, T. C. & Hore, P. Viability of superoxide-containing radical pairs as magnetoreceptors. The Journal of chemical physics 151 (2019).

35. Husen, P., Nielsen, C., Martino, C. F. & Solov’yov, I. A. Molecular oxygen binding in the mitochondrial electron transfer flavoprotein. Journal of Chemical Information and Modeling 59, 4868–4879 (2019).

36. Pottosin, I. & Zepeda-Jazo, I. Powering the plasma membrane Ca2+-ROS self-amplifying loop. Journal of Experimental Botany 69, 3317–3320 (2018).

37. Chambers, J. W. & LoGrasso, P. V. Mitochondrial c-Jun N-terminal kinase (JNK) signaling initiates physiological changes resulting in amplification of reactive oxygen species generation. Journal of Biological Chemistry 286, 16052–16062 (2011).

38. Dikalov, S. Cross talk between mitochondria and NADPH oxidases. Free Radical Biology and Medicine 51, 1289–1301 (2011).

39. Tiwari, Y., Raghuvanshi, P. & Poonia, V. S. Radical pair mechanism and the role of chirality-induced spin selectivity during planaria regeneration. Applied Physics Letters 125, 103701. issn: 0003-6951. eprint: https://pubs.aip.org/aip/apl/article-pdf/doi/10.1063/5.0227302/20142649/103701\_1\_5.0227302.pdf. 10.1063/5.0227302 (Sept. 2024).

40. Haberkorn, R. Density matrix description of spin-selective radical pair reactions. Molecular Physics 32, 1491–1493 (1976).

41. Rodgers, C. T., Norman, S. A., Henbest, K. B., Timmel, C. R. & Hore, P. Determination of radical re-encounter probability distributions from magnetic field effects on reaction yields. Journal of the American Chemical Society 129, 6746–6755 (2007).

42. Solov’yov, I. A. & Schulten, K. Magnetoreception through cryptochrome may involve superoxide. Biophysical journal 96, 4804–4813 (2009).

43. Evans, E. W. et al. Sub-millitesla magnetic field effects on the recombination reaction of flavin and ascorbic acid radicals. The Journal of Chemical Physics 145 (2016).

44. Kattnig, D. R. et al. Chemical amplification of magnetic field effects relevant to avian magnetoreception. Nature Chemistry 8, 384–391 (2016).

45. Kattnig, D. R., Sowa, J. K., Solov’yov, I. A. & Hore, P. Electron spin relaxation can enhance the performance of a cryptochrome-based magnetic compass sensor. New Journal of Physics 18, 063007 (2016).

46. Player, T. C. & Hore, P. Source of magnetic field effects on the electrocatalytic reduction of CO2. The Journal of Chemical Physics 153 (2020).

47. Inc., W. R. Mathematica, Version 13.1 11.0 Champaign, IL. https://www.wolfram.com/mathematica.

48. Vučković, J. et al. Construction and Application of a Static Magnetic Field Exposure Apparatus for Biological Research in Aqueous Model Systems and Cell Culture. Bioprotocol 14, e5077 (2024).

